# A multiscale model predicts the sensitivity of *Chlorella vulgaris* to light and nitrogen levels in photobioreactors

**DOI:** 10.1101/2021.04.14.439858

**Authors:** Juan D. Tibocha-Bonilla, Cristal Zuniga, Jared T. Broddrick, Karsten Zengler, Rubén D. Godoy-Silva

## Abstract

The maximization of lipid productivity in microalgae is crucial for the biofuel industry, and it can be achieved by manipulating their metabolism. However, little efforts have been made to apply metabolic models in a dynamic framework to predict possible outcomes to scenarios observed at an industrial scale. Here, we present a dynamic framework for the simulation of large-scale photobioreactors. The framework was generated by merging the genome-scale metabolic model of *Chlorella vulgaris* (*i*CZ843) with reactor-scale parameters, thus yielding a multiscale model. This multiscale model was employed to predict the sensitivity of growth and composition variation of *C. vulgaris* on light and nitrogen levels. Simulations of lipid accumulation quantified the tradeoff between growth and lipid biosynthesis under nitrogen limitation. Moreover, our modeling approach quantitatively predicted the dependence of microalgal metabolism on light intensity and circadian oscillations. Finally, we use the model to design a reactor irradiance profile that maximized lipid accumulation, thus achieving a lipid productivity increase of 46% at a constant intensity of 966 μE m^−2^ s^−1^. Our modeling framework elucidated how metabolism and external factors can be combined to predict optimized parameters for industrial applications.

## Background

Microalgae are unicellular photosynthetic organisms that fix carbon dioxide (CO2) in presence of light to obtain energy and synthesize necessary metabolic precursors for growth. Carbon fixation of microalgae can be up to ten times higher than that of plants [1,2] and accounts for about 40% of the Earth’s CO2 fixation [3]. Their powerful photosynthetic activity caused microalgae to be a focus of research in ecology, systems biology, and bioengineering.

Oleaginous microalgae are able to store lipids at levels higher than 20% w/w, which is the minimum threshold for profitable production of biofuels [4,5]. *Chlorella vulgaris* has drawn widespread attention, as it can intracellularly concentrate up to 50% w/w of lipids in form of triacylglycerols (TAGs) [6]. Since TAGs serve as main precursors for biofuel production, *C. vulgaris* is a promising lipid producer with potential application at industrial scale. However, studies about lipid accumulation have shown that stress conditions trigger lipids accumulation at the expense of decreased growth rate [6,7]. At the industrial scale, reactor lipid productivities are severely limited by this tradeoff, rendering the study of the interwoven connection among metabolic and physical drivers of lipid biosynthesis as a field of great significance and research focus. Different efforts have been made towards manipulating microalgae metabolism, mainly by varying light, nitrogen, and growth conditions [8]. Nonetheless, the maximization of lipid productivity in microalgae has remained a challenge for years, due to experimentation being extremely time- and resource-intensive. Therefore, computational tools appear as a promising alternative, since they can assess optimal growth conditions time- and cost-effectively.

To date, all but one study on dynamic modeling of the growth of *C. vulgaris* have been based on kinetic modeling [2,9–17]. All kinetic models are based on a black-box framework. That is, their primary focus is fitting experimental data regardless of the structure of underlying phenomena. These models have been crucial for the development of the chemical and biochemical industry [18]; however, a more robust approach is necessary to predict not only global reactor dynamics, but also intracellular metabolic capabilities, namely lipid biosynthesis.

Microalgal metabolism has been previously studied using genome-scale metabolic (GSM) models, thus elucidating organelle functionality and pathway coupling, as well as the interactions of photosynthetic pathways with the rest of the network under different light wavelengths [19,20]. Some studies on microalgae metabolic modeling have incorporated photon uptake [21,22], enabling process optimization of light intensity and culture density at the laboratory scale [21]. In recent studies, experimental time-course biomass compositions were incorporated in the GSM model to predict the metabolic response to nitrogen depletion [23] and to optimize nitrate supply in *C. vulgaris* [24]. Notably, GSM modeling allowed to assess metabolic crosstalk in a *C.vulgaris and Sacharomyces cerevisiae* synthetic syntrophic community [25]. Though, there exists no mathematical framework that combines these metabolic models with reactor-scale dynamics for the prediction of growth of microalgae, such as *C. vulgaris*, at a scale relevant to industrial applications.

Merging mathematical representations of phenomena at the genome and reactor scale would result in a multiscale model. Separately, each mathematical representation is successful in modeling the phenomenon they were based upon but cannot individually account for entire reactor dynamics. For example, a genome-scale metabolic (GSM) network is a powerful tool for understanding species-specific metabolism. However, without experimental input, a GSM model cannot predict time-course metabolic changes, as well as reactor-scale growth.

Here, we generated a multiscale model that simulates time-course growth and biomass composition variations using an available GSM model of *C. vulgaris (i*CZ843). The model includes light attenuation and uptake, photoinhibition, nitrogen and carbon uptake kinetics, and carbon allocation (carbohydrate and lipid accumulation and consumption). We then employed previously reported experimental growth data under different nitrogen and light conditions to validate our predictions.

For the former, we employed data by Adesanya et al. [13] as reference, whereas for the latter we used data by Kim et al. [14]. As different strains can have different metabolic capabilities, we performed data regression and validation individually for both cases. Finally, we present the model’s potential for its use in reactor design and optimization by determining light profiles for maximizing lipid productivity in a photobioreactor.

## Results

### Prediction of growth trends at two different initial nitrogen concentrations

So far, dynamic modeling of the growth of *C. vulgaris* has been almost entirely addressed using kinetic modeling (Table 1). However, this type of modeling lacks the ability to include the effect of reactor-scale dynamics on microalgal metabolism. Moreover, no model has accounted for the combined effect of nutrient uptake, circadian oscillations, and light attenuation on the dynamic growth and biomass composition of microalgae.

**Table 1.** Existing models for the growth of *Chlorella vulgaris*. Early models focused on dynamic modeling through black-box methods.

| Study (Year) | Model characteristics* |  |  |  |  |  |  |  |  |
| --- | --- | --- | --- | --- | --- | --- | --- | --- | --- |
|  | Type | D | CU | NU | LD | LU | CO | LA | SA |
| Iehana (1990) [9] | K | ✓ |  |  |  |  |  |  |  |
| Wijanarko et al. (2004) [10] | K | ✓ | ✓ |  |  |  |  |  |  |
| Filali et al. (2011) [11] | K | ✓ | ✓ |  |  | ✓ |  |  |  |
| Concas et al. (2013) [12] | K | ✓ | ✓ |  | ✓ | ✓ | ✓ |  |  |
| Adesanya et al. (2014) [13] | K | ✓ | ✓ | ✓ | ✓ | ✓ |  | ✓ |  |
| Kim et al. (2015) [14] | K | ✓ |  |  | ✓ | ✓ |  |  |  |
| Sakarika et al. (2016) [15] | K | ✓ | ✓ |  |  |  |  | ✓ |  |
| Adamczyk et al. (2016) [2] | K | ✓ | ✓ |  |  |  |  |  |  |
| Chan et al. (2016) [16] | K | ✓ | ✓ |  |  | ✓ |  |  |  |
| Zuñiga et al. (2016) [20] | GS |  | ✓ | ✓ |  | ✓ |  |  |  |
| Zuñiga et al. (2017) [6] | GS | ✓ | ✓ | ✓ |  | ✓ |  |  |  |
| Mansouri et al. (2017) [17] | K | ✓ |  |  |  |  |  |  |  |
| Chien-Ting Li (2019) | GS | ✓ | ✓ | ✓ | ✓ |  |  | ✓ |  |
| Tibocha-Bonilla et al. (2020, this study) | MS | ✓ | ✓ | ✓ | ✓ | ✓ | ✓ | ✓ | ✓ |

\* Model characteristics are abbreviated as follows: dynamic (D), carbon uptake (CU), nitrogen uptake (NU), light distribution (LD), light uptake (LU), circadian oscillations (CO), lipid accumulation (LA) and starch accumulation (SA). Models were classified into three types: kinetic (K), genome-scale (GS) and multiscale (MS).

We generate a multiscale model capable of predicting this interaction by merging different mathematical representations (or modules, see Fig 1A) of biological and reactor dynamics at different scales. Nitrogen availability has been identified as one of the most important drivers of microalgal growth, as it has a profound impact on the cellular phenotype of phototrophs [6]. To validate our model’s sensitivity to varying nitrogen concentration, we contrasted predictions of our model to experimental data reported in a kinetic study by Adesanya et al., in which *C. vulgaris* was cultivated under two different initial nitrogen concentrations (first scenario: 0.021 g L^−1^, second scenario: 0.124 g L^−1^) and macromolecular cellular contents were recorded [13] (markers in Fig 1B-E).

**Fig 1.**
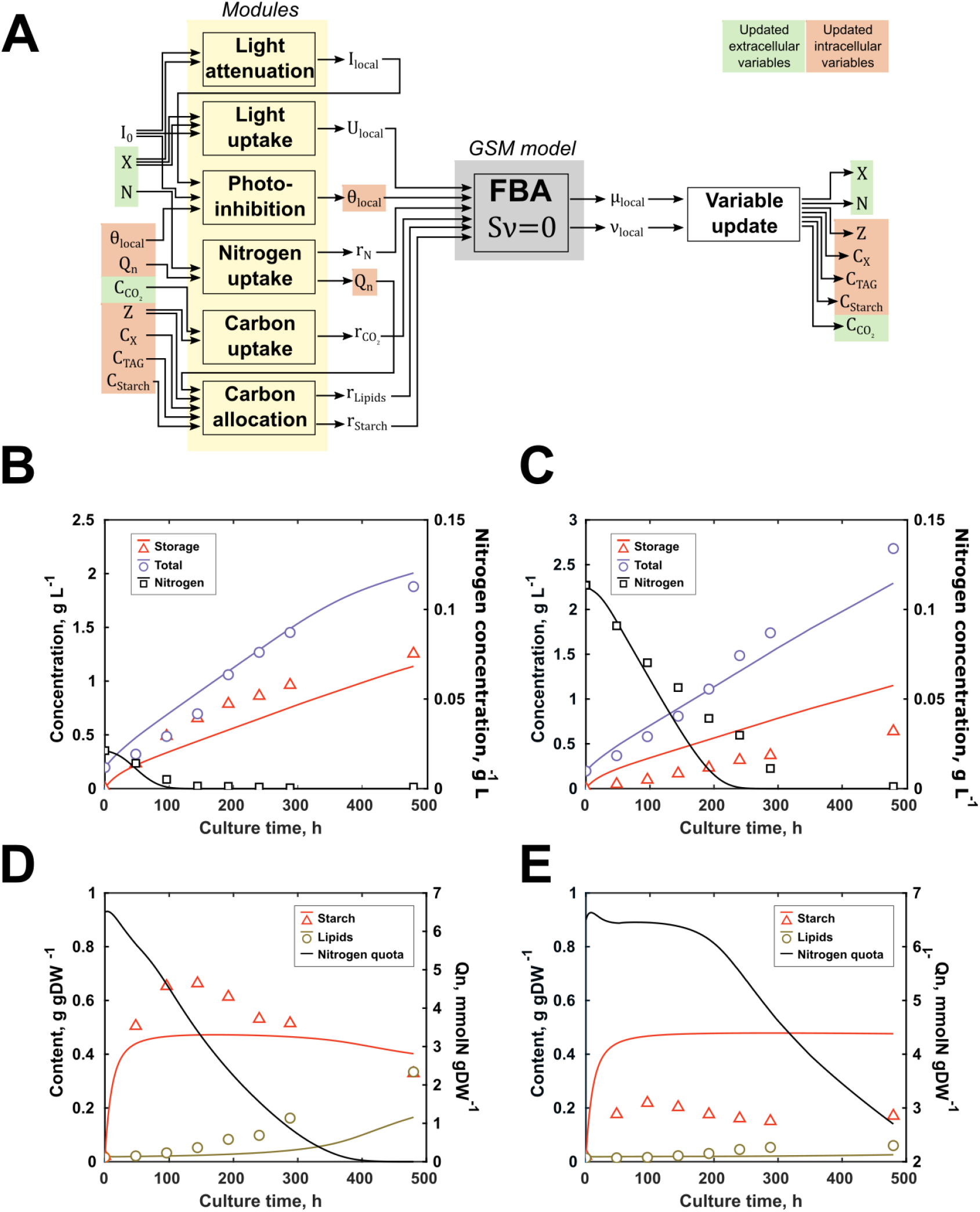
Simulated and reported data of *Chlorella vulgaris* at two different initial nitrogen concentrations. (A) Schematic representation of the numerical algorithm employed in this work for a single timestep and light interval (see *Methods*). (B) and (D) correspond to the experiment with an initial nitrogen concentration of 0.021 g L^−1^, while (C) and (E) were recorded under one of 0.124 g L^−1^. All experiments were reported and simulated under a continuous irradiance of 80 μE m^−2^ s^−1^. Continuous lines and markers represent predicted data by our model and reported data by Adesanya et al. [13], respectively.

We used the first scenario, with an initial nitrogen concentration of 0.021 g L^−1^ (Fig 1B & D) for the calculation of strain-specific parameters (shown in Table S1) and simulated a second scenario (Fig 1C & E) to test for predictive capability. In this scenario, a relatively low initial nitrogen concentration in the media (half that of standard BBM medium [26]) caused the size of the internal nitrogen pool to decrease steadily throughout the culture duration (Fig 1D). Since the microalga was not able to replenish its nitrogen reserves, lipid accumulation was triggered 100 h after nitrogen was depleted from the medium.

As shown in Fig 1C, under the growth conditions of the second scenario, nitrogen availability was increased six-fold. Though there were some quantitative differences regarding the starch content of *C. vulgaris*, our model was able to capture the overall trends. Our simulations show that a significant increase in the nitrogen availability allowed the microalga to replenish its nitrogen reserves for the first 200 h and caused it to deplete nitrogen from the medium 130 h later than the first scenario (Fig 1E).

The model was able to capture the almost complete inactivation of the lipid metabolism due to this nitrogen availability. However, despite there being almost no lipid accumulation and high nitrogen levels, the predicted growth rate was only amplified from an average of 0.0032 to 0.0038 h^−1^ (19% increase), similar to an experimentally determined increase of 15%.

### Simulation at different light intensities

We accounted for spatial light distributions, as energy metabolism is sharply hindered for cells further away from the light source, especially in larger-scale vessels. In addition, we included the modeling of photoinhibition, since it restricts the maximum amount of light a culture can be subjected to and the duration of exposure. Although several species have been shown to adapt to high light conditions in the long term [27,28], photoinhibition still significantly diminishes the growth capability of phototrophs [29–31], and specifically of *C. vulgaris* above 2400 μE m^−2^ s^−1^ [32]. According to previous reports, electron transfer through the photosystems is controlled by the fraction of active protein D1 of the photosystem II (PSII). Therefore, we used the model proposed by Baroli et al. [30] to determine the fraction of active D1 protein as a means to penalize the effective photon input to the metabolic network.

We employed previously reported data to validate the model’s sensitivity to change in light intensities. Kim et al. [14] cultured *C. vulgaris* at six different irradiances and monitored biomass concentration throughout the timespan of the culture. We used data recorded at 848 μE m^−2^ s^−1^ for the regression of parameters (Fig 2), while data at 30, 55, 80, 197, and 476 μE m^−2^ s^−1^ were employed for model validation.

**Fig 2.**
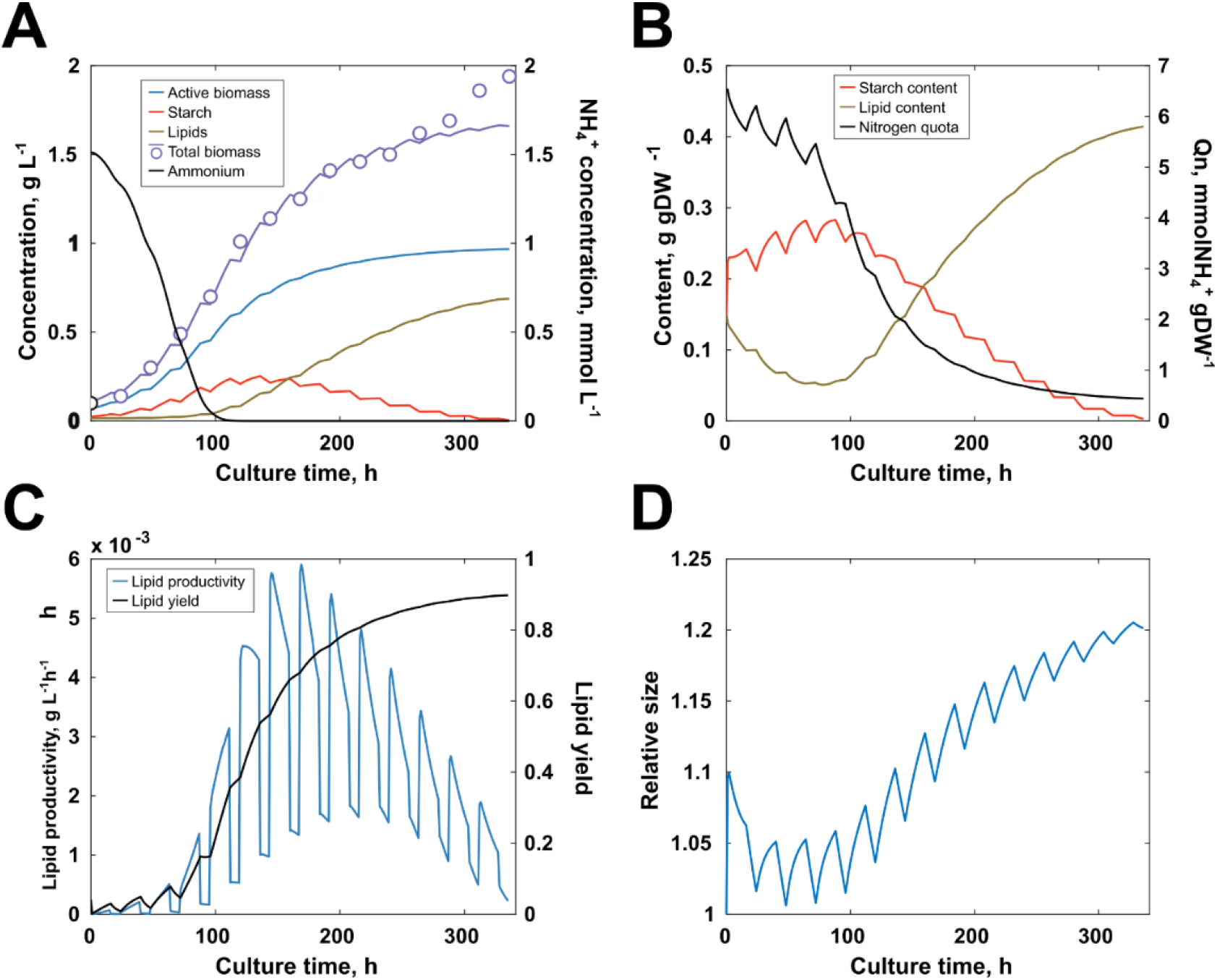
Contrast of calculations and reported data at 848 μE m^−2^ s^−1^ by Kim et al [14]. (A) Global reactor concentrations of active (non-storage) biomass, starch, lipids, total biomass and nitrate contrasted with reported data of total biomass. (B) Intracellular content of starch, lipids and nitrogen. (C) Contrast of lipid productivity with lipid yield (% of carbon input directed to lipid production). (D) Variation of cell size. Continuous lines and markers represent predicted data by our model and reported data, respectively.

Experimental total biomass concentrations were recapitulated by our model. Even though intracellular concentrations were not measured by Kim et al. [14], the model could be used to hypothesize microscopic and macroscopic phenomena that lay underneath, e.g. circadian clock oscillations, carbon allocation and responses to nitrogen depletion. As opposed to the case of Adesanya et al. [13], our simulations showed that an elevated irradiance of 848 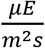 made impossible for the microalga to maintain the intracellular nitrogen levels after the nitrogen was depleted from the medium at 100 h. This caused the lipid production to be triggered around the same time point, as intracellular nitrogen levels were already decreasing sharply (Fig 2B).

According to our simulations, the optimum lipid productivity was achieved shortly after nitrogen was consumed from the medium but rapidly decreased afterward (Fig 2C). Interestingly, even though nitrogen depletion from the medium was achieved at 100 h, peak global lipid productivity of the photobioreactor took place at 168 h, when internal reserves were running low but were not yet depleted. Overall lipid productivity decreased steadily for the following 167 h, as growth was increasingly hindered by internal nitrogen depletion and the cell size was reaching its limit.

Our simulations predicted cell size change as a result of light/dark cycles and long-term nitrogen depletion (Fig 2D). In general, during the light period, the cell focuses on the accumulation of starch for later use under dark conditions, causing its size to increase. The opposite behavior occurs during the dark period, in which starch is consumed for maintenance and growth. In the long term, nitrogen depletion-induced lipid accumulation yielded bigger cells at the late stage.

Simulations successfully reproduced growth behavior in the photobioreactor at different irradiance conditions (Fig 3). Even though they underestimated biomass production at low intensities, the overall growth trends were predicted accurately. Since shading hampers the ability to fix inorganic carbon, the slope of the growth curves decreased with lower irradiances. At higher intensities, it is evident that an increase from 476 to 848 μE m^−2^ s^−1^ did not signify an improvement in the overall culture growth. Our simulations show that this was a consequence of a combined effect of shading, photoinhibition and nitrogen limitation.

**Fig 3.**
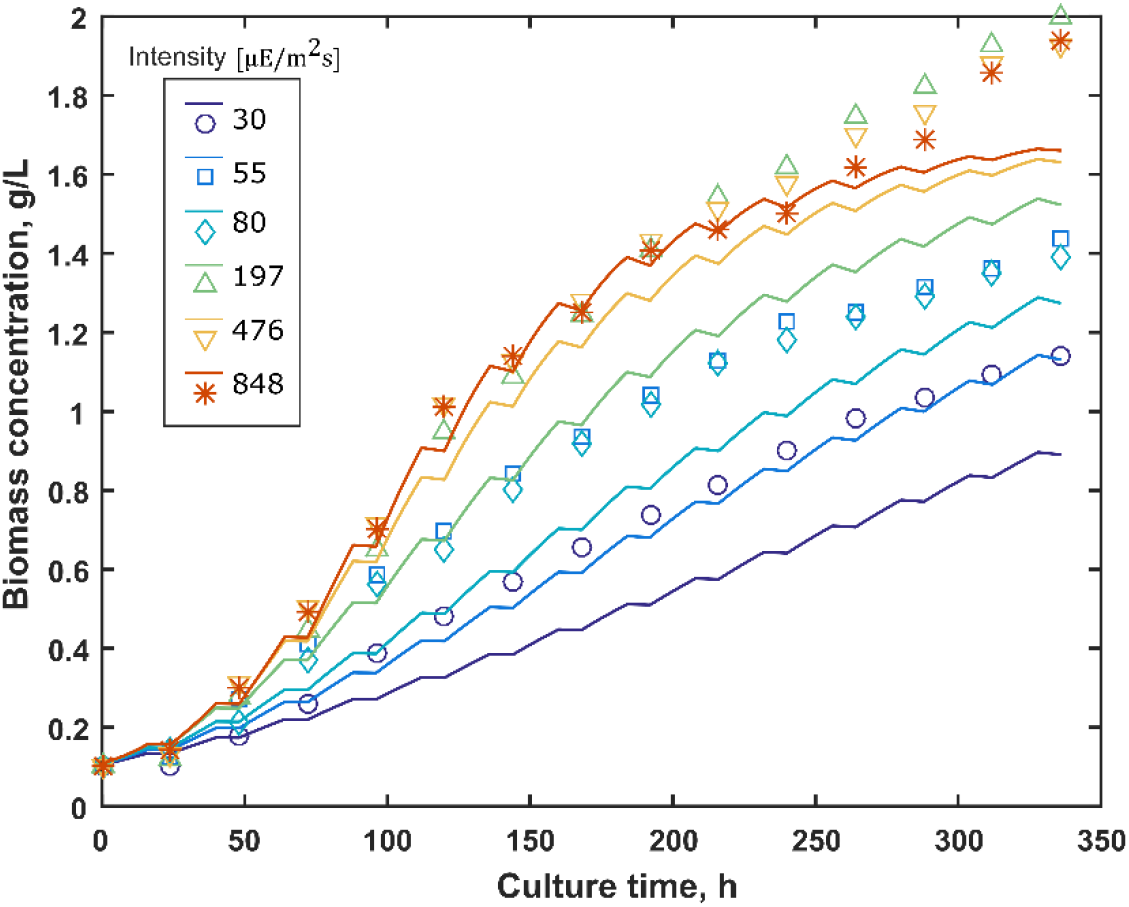
Growth simulation and experimental validation for different irradiance conditions reported by Kim et al. [14] at. Lines represent model simulations while markers show reported experimental data. Even though the model underestimated growth at low intensities, growth rate response to varying light intensities was captured by the model.

### Optimization of the lipid productivity in a photobioreactor

In order to illustrate the model’s aptitude for process design and optimization, we predicted the optimal light strategy to maximize lipid productivity in a stirred tank photobioreactor reactor with six internal radially distributed fluorescent lamps.

In brief, five variables were manipulated to search for the optimal global lipid productivity condition: lamp irradiance at time zero I_0_, lamp irradiance at the end of the culture I_f_, photoperiod p, culture duration t_f_, and shape of irradiance temporal profile (see *Methods*) represented by the coefficient b_I_. A hypothetical base case was given to the model as the initial condition of the optimization, with an irradiance profile within the typical values previously used for experimental optimization of lipid accumulation [29]. Fig S3 shows a summary of the simulation results of the base case, and Table 2 summarizes the base case and optimized values of the manipulated variables.

**Table 2.** Initial and final values of the manipulated variables of the optimization.

| | $I_0$<br>$\mu\text{E m}^{-2} \text{ s}^{-1}$ | $I_f$<br>$\mu\text{E m}^{-2} \text{ s}^{-1}$ | $p$<br>$h$ | $t_f$<br>$h$ | $b_l$ |
| --- | --- | --- | --- | --- | --- |
| Base case | 400 | 600 | 16 | 300 | 0.5 |
| Optimized | 966 | 966 | 17 | 374 | 0.0 |

Our model predicted an optimum constant irradiance of 966 μE m^−2^ s^−1^. As shown in Fig 4, this light intensity improves the final biomass concentration from 1.70 to 1.83 g/L in 374 h, which is roughly a 7% increase in biomass, and 46% increase in final lipid concentration, for a 25% longer culture timespan.

**Fig 4.**
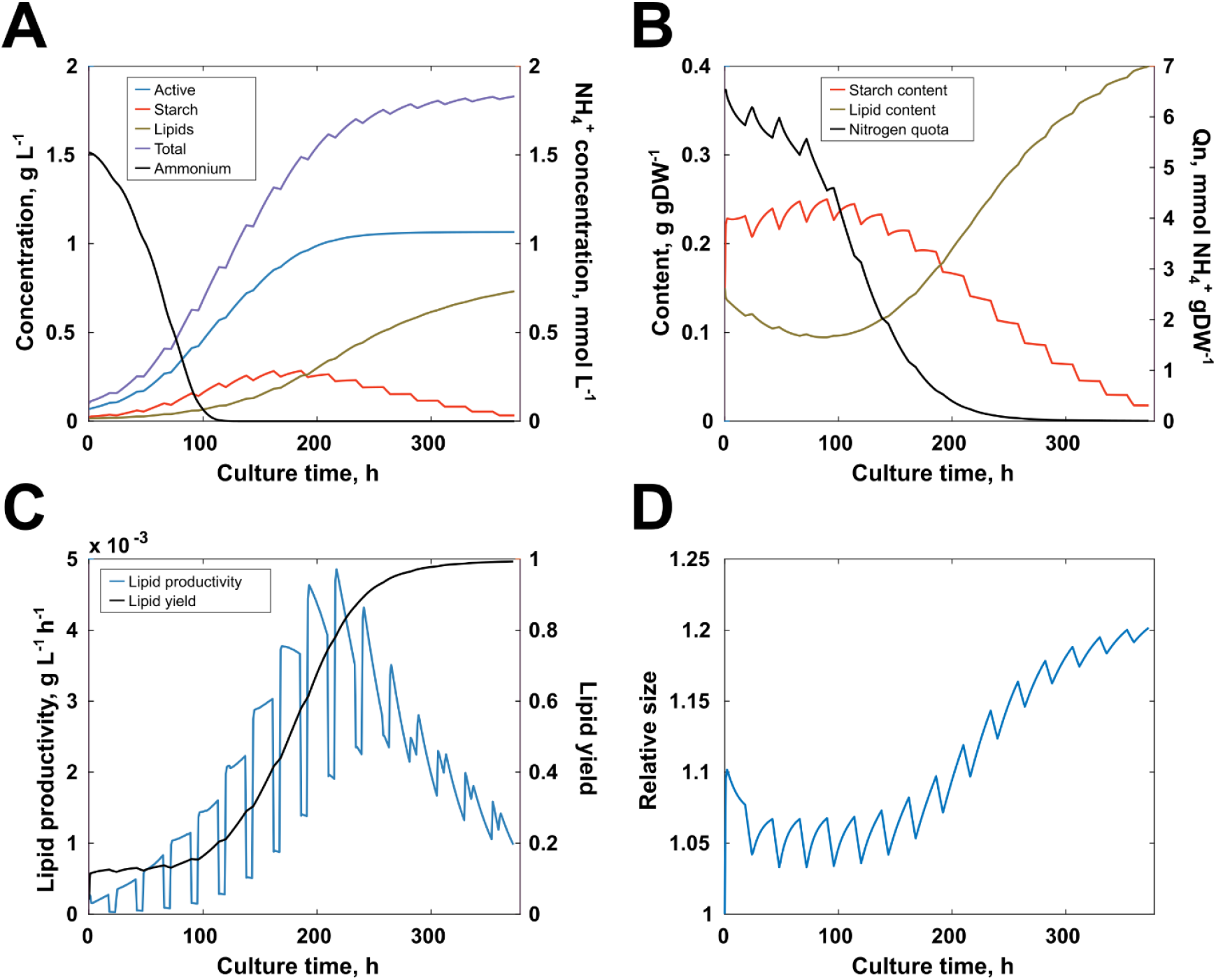
Simulation results of lipid productivity maximization by varying light strategy, photoperiod and culture time. (A) Resulting optimal global reactor concentrations of active biomass, starch, lipids, total biomass and nitrate contrasted. (B) Intracellular content of starch, lipids and nitrogen. (C) Lipid productivity and lipid yield (% of carbon input directed to lipid production). (D) Variation of cell size.

Even though at the early stage of (0 – 100 h) a fraction of the culture is subjected to an irradiance of around 3,000 μE m^−2^ s^−1^ (Fig 4), a large portion of it is under a much lower but still significant irradiance of 200 μE m^−2^ s^−1^. This, along with high nitrogen availability, favored higher growth rates in such a way that photoinhibition was compensated. Moreover, the optimization showed that a photoperiod of 17:7 is sufficient to satisfy dark period metabolic requirements without negatively affecting growth or lipid productivity.

As early as 25 h into the culture, the highest irradiance inside the culture lowers to 2400 μE m^−2^ s^−1^. During the medium-growth stage (100 - 200 h) shading rapidly diminished photoinhibition from approximately 80% to 25%, as highest irradiances were of only 800 μE m^−2^ s^−1^, and the average dropped to 200 μE m^−2^ s^−1^.

At the low-growth stage (200+ h), shading is so substantial that the average irradiance drops to 142 μE m^−2^ s^−1^ and stabilizes there for the rest of the culture. Moreover, as exhibited in Fig 5, at this point light uptake had almost halted throughout the majority of the reactor, with an average photon uptake of only 70 mmol gDW^−1^ h^−1^, as opposed to an average of 1000 mmol gDW^−1^ h^−1^ at the high-growth stage.

**Fig 5.**
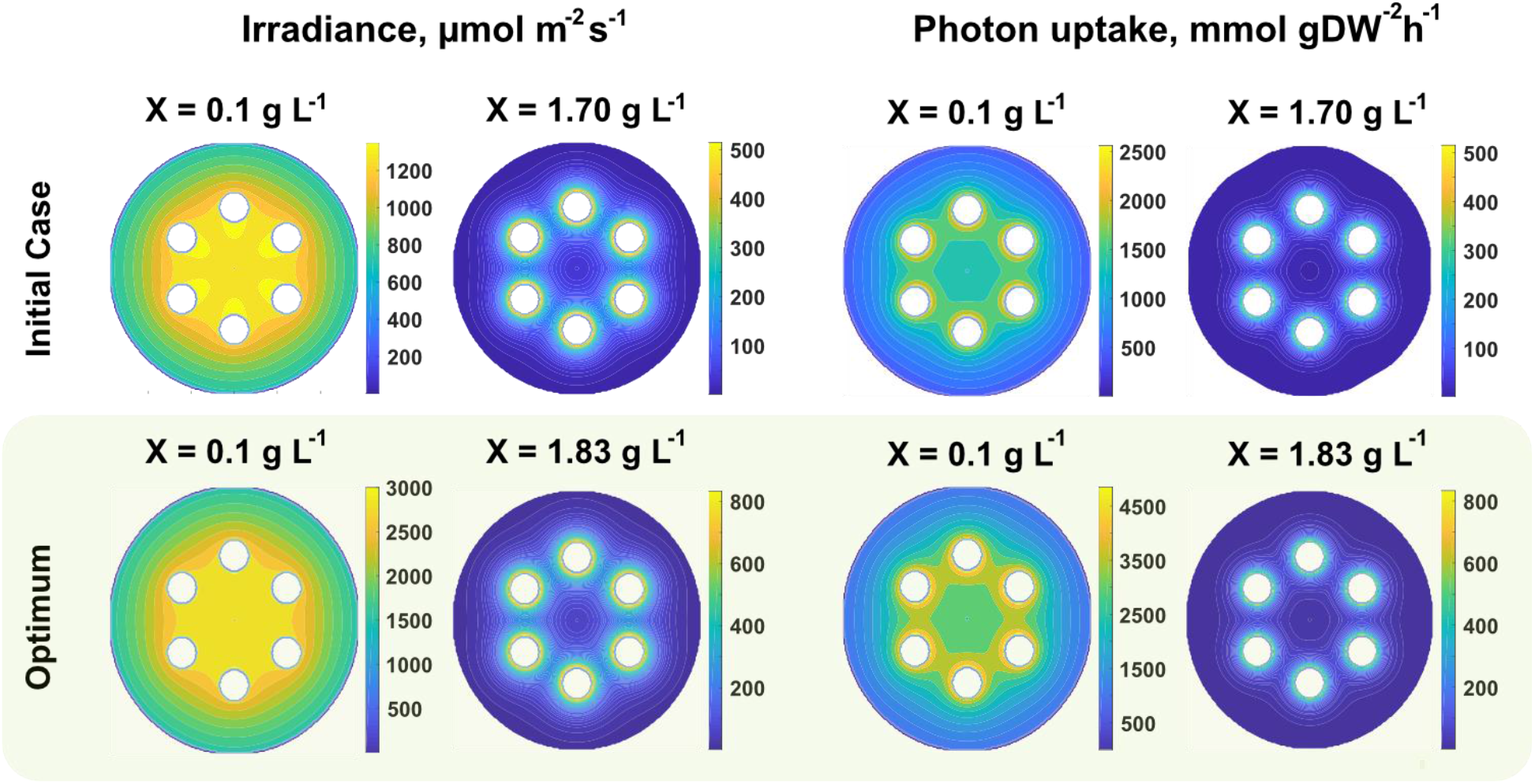
Light and photon uptake distributions at the beginning and end of the culture. Internal cross-section distributions in the bioreactor are shown for the initial and optimum cases, at the initial and final concentration. Distributions shown correspond to light intensity (irradiance) and photon uptake. (Top row) Distributions for the initial case. (Bottom row) Distributions for the found optimum condition.

## Discussion

### The multiscale model accurately predicts dynamic growth and biomass breakdown

In this work, we generated a multiscale metabolic model that predicts the growth dynamics of *Chlorella vulgaris*, by coupling mathematical representations for circadian oscillations, substrate uptake, photoinhibition and light uptake distributions. Kinetics-based dynamic metabolic modeling has already been conceived [33,34], but only Jeong et al. [35] have applied it on a complete metabolic network, and one study on a sub-network [36]. Moreover, neither study accounted for time-dependent carbon allocation or light uptake distributions.

First, we used the reported growth kinetics by Adesanya et al. [13] at two different initial nitrogen concentrations to test the model’s ability to capture differential reactor behavior when nitrogen availability changed. In the first scenario, the microalga was subjected to a relatively low nitrogen availability, thus reducing its uptake rate, and preventing it from replenishing its nitrogen reserves. Consequently, lipid accumulation was triggered after nitrogen was depleted in the medium; however, its activation was not significant until days later. This implies that nitrogen depletion from the medium signifies the beginning of the end of exponential growth, rather than the end itself. A similar behavior was obtained by Mansouri et al. [17] under a comparable setup. They reported that exponential growth was maintained for the first 96 h of growth, time after which growth gradually stopped until their last recorded instance at 168 h.

A second scenario was tested in which the microalga was cultured with a six-fold increase in initial nitrogen concentration. As expected, lipid accumulation dramatically decreased in both their report and our predictions, but growth did not increase significantly (19 %). Experimental and predicted growth rates remained one order of magnitude lower than previously reported maximum growth rates of 0.039 [6] and 0.033 h^−1^ [14,17]. Our simulations show that this was mainly caused by the comparatively reduced irradiance as opposed to common working irradiances of 648 [6,20], 20 – 1400 [37], and 30 – 848 [14] μE m^−2^ s^−1^. Low light irradiance hindered the overall growth rate of the microalga, which, in addition to its higher nitrogen availability, allowed it to replenish its nitrogen reserves for the most part of the culture. This relatively nitrogen-replete condition caused storage molecule (lipid and starch) production to drop and rendered lipid accumulation almost non-present.

A visible over-estimation of starch content in the second scenario was mainly caused by the prioritization of starch consumption in the dark period of the topology of our carbon allocation algorithm (Fig S2), which induces error when trying to predict a permanently illuminated culture, as used by Adesanya et al. [13]. Further work on the generation of a multiscale model for the mixotrophic growth of *C. vulgaris* will be necessary for this model to properly include starch consumption in the light period, with quantitative accounting of carbon allocation and the differential destination of carbon with concomitant starch breakdown and carbon dioxide consumption.

### Light intensity drives oscillations in biomass components and cell size

As a next step, we used reported data by Kim et al. [14] to show the model’s capability to predict the effect of varying the light intensity. Our model was able to reproduce the experimental data and gave insight into the phenomena that caused the growth trends. For example, circadian clock oscillations are evident in all monitored variables. Fig 2B shows the starch accumulation-consumption cycles, along with a macroscopic interchange of carbon flow between starch and lipids after nitrogen depletion. This behavior has previously been quantitatively determined for other oleaginous microalgae, such as *Synechococcus elongatus* [38], *Chlorella sorokoniana* [39] and *Dunalliela salina* [40]. In one study, *S. elongatus* exhibited a peak in ADP-glucose pyrophosphorylase activity, as well as in glycerol-3-phosphate (G3P) production from ribulose biphosphate (RuBP) coming from the reductive pentose-phosphate pathway (PPP) close to dawn, implying high starch production in the light period [38]. The same study found the exact opposite behavior in the dark, with peak activities of glycogen debranching enzyme (glgX).

The light-induced storage-consumption cycle is visible in the cell size variations through time, where cells can be expected to increase in size in the light while storing starch and do the opposite in the dark, as previously reported by Martins et al. for the cyanobacterium *Synechococcus elongatus* [41]. Furthermore, our simulations captured the well-known tradeoff behavior between specific lipid biosynthesis and growth rate, which causes the long-term cell size increase after nitrogen depletion.

### The model can be used to design light strategies for increased lipid productivity

We used the model to maximize the lipid productivity in a case of study to illustrate its potential for reactor design and optimization. As opposed to previous trials on light strategies, in which irradiance increases in a stair-step fashion [29], the model predicted an optimal global light productivity at a constant lamp irradiance of 966 μE m^−2^ s^−1^. At this irradiance lipid productivity was predicted to increase in 46 %, despite global biomass concentration increasing by only 7%.

Even though the optimal irradiance is relatively high, extreme values are only reached at the early stage of culture, during which high nitrogen availability favored higher growth rates in such a way that photoinhibition was compensated. Moreover, during the first day of culture the highest irradiance inside the culture lowered to 2400 μE m ^−2^ s ^−1^, which, according to Pfendler et al. [32], is the limit above which photoinhibition seriously reduces light uptake and growth in *C. vulgaris*.

At the late stage, shading protects most of the cells from excessive irradiances and keeps it at a level that still favors metabolic activity. At this point, increasing the irradiance of the lamps would hardly alter the global light availability and uptake, but would critically increase irradiance in the vicinity of the lamps, where nitrogen-deplete microalgae are no longer capable of compensating for photoinhibition. This means that, for this case, an individual lamp irradiance of 966 μE m^−2^ s^−1^ is high enough to boost growth without excessively hindering photon uptake at all culture stages. As a result, this irradiance is optimal for overall growth and lipid production in the photobioreactor.

In addition, it is worth noting that our model does not yet include neither the modeling of heat transfer mechanisms between the lamps, medium and surroundings, nor mixing phenomena which causes it to assume every cell is subjected to the same temperature. With this, the suggested optimum is only attainable under a cooling system that is efficient enough to maintain overall temperature between 22 – 26 °C [42].

## Conclusions

Modeling of microalgal growth is key for the industrial production of biofuels, as it allows for the design and optimization at a reactor- and whole plant-level. For this purpose, it is necessary to generate a framework including accurate accounting of phenomena at every relevant scale, namely at the genome scale and reactor scale. In this work, combined a GSM network with mathematical representations of circadian oscillations, nutrient uptake and light distributions, which yielded a comprehensive multiscale model of the growth dynamics of *Chlorella vulgaris*. We included a detailed framework for the calculation of photon uptake at different irradiances, by considering light attenuation and photoinhibition. The model was tested both at different nitrogen levels and light irradiances, rendering it capable of being used for the prediction of specific conditions that maximize lipid productivity.

## Methods

All simulations were carried out within the MATLAB 2016b (MathWorks Inc.) environment and using the COBRA Toolbox v3.0 [43]. Dynamic Flux Balance Analysis (dFBA) was used for time-course flux distribution calculations and concentration updates, and GUROBI 7.5.2 was employed as the solver for the linear optimization problems. A more detailed explanation of the model’s algorithms is shown in this section.

### The multiscale metabolic model

At the core of our calculations lies the genome-scale metabolic model of the oleaginous microalga *Chlorella vulgaris: iCZ843* [20], with previously proposed modifications for both heterotrophic and autotrophic growth [6,20]. Overall, the GSM was solved using COBRA Toolbox (dFBA) for metabolic flux distributions. In addition, a set of additional models were included to account for secondary phenomena which constrained the solution space of the Linear Programming (LP) system (GSM in Fig 1A). Phenomena were included according to previous reports of specific physical and physiological mechanisms significantly affecting growth. Included mechanisms were light attenuation, light uptake, photoinhibition, nitrogen and carbon uptake kinetics, and carbon allocation (carbohydrate and lipid accumulation and consumption). Mixing, heat and mass transfer phenomena were not included in the present model. A simplified representation of the general numerical algorithm is presented in Fig 1A.

### Light attenuation

Several studies have focused on light attenuation of microalgae [44–49], with a few solely on *Chlorella vulgaris* [44,45]. In this work, we decided to use the model for light absorption and scattering proposed by Naderi et al. [45], which enabled accurate predictions of light distribution at low and high cell densities. The intensity profile function is shown in Eq. (1).

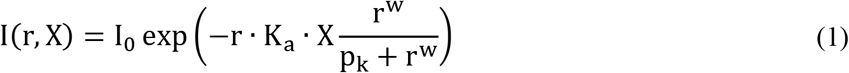

For internally illuminated reactors, the distance *r* was computed as the distance between the edge of the light source and any given point inside the culture. Several internal sources were accounted for by taking the sum of the calculated individual light distributions. For externally illuminated (jacketed) reactors *r* was calculated as the distance between the illuminated edge of the reactor and any given point inside the culture. K_a_ represents the absorption coefficient and is a function of the biomass concentration X and the maximum absorption coefficient K_a,max_, as shown in Eq. (2). In addition, p_k_, b and w are model parameters.

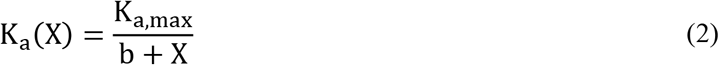

The initial intensity *I*_0_ corresponds to either the nominal or measured intensity of the light source, whichever was reported in the studies. Absorption and scattering coefficients were left unchanged throughout the culture time, although further studies can compute time-specific coefficients from absorption spectrum data [21].

For increased computation speed, we divided the photobioreactor in a finite number of zones with the same light intensity and calculated the overall reaction rates as a volume-weighted average of the individual intervals. The number of light intervals (N_I_) were determined in a logarithmic fashion, as shown in Eq. (3). Mesh dependence analyses showed that 10 active (with non-zero irradiance) intervals were enough for the simulations to be independent of the number of intervals.

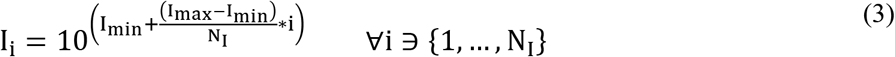

### Light uptake

We defined a photon conservation balance over a differential element (Fig S1) to account for spatial light uptake distribution, as shown in Eq. (4).

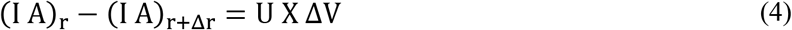

The conservation balance is readily converted to the differential equation shown in Eq. (5), and a cellular uptake profile (U) is obtained in Eq. (6). The magnitude U is at this point a unit-consistent input to the GSM model of the microalga, which represents the upper bound of specific photon uptake rate.

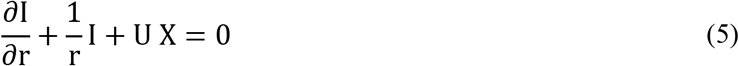

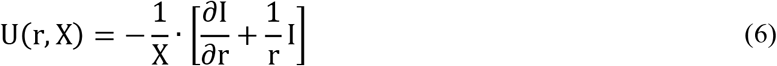

A similar procedure for a planar reactor yields the homologous expression shown in Eq. (7).

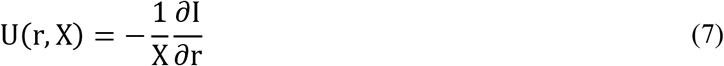

### Photoinhibition

Photoinhibition is the reduction of photosynthetic capacity in photoautrotrophic organisms [50], and has been proven to be controlled by the photodamage – repair dynamics of the protein D1 of the photosystem II (PSII) [31,50]. Consequently, an accurate account of the fraction of available photons that reach the metabolic network can be performed by determining the fraction of active protein D1. Therefore, the effect of photoinhibition in the model was represented by the fraction of active D1, employing the model by Han [50], with the coefficients reported by Baroli et al. [30] (see Eqs. (8) and (9)). The magnitude θ represented the fraction of photons that were used by the metabolic network, and is a function of time t, the first-order D1 photorepair and photodamage coefficients, k_r_ and k_d_, and light intensity I. Moreover, k_r_ is a function of light intensity, which follows a linear behavior described by the slope m_k_ and intersect b_k_.

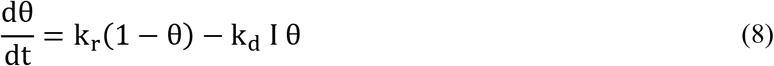

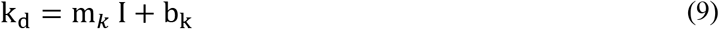

At every timestep and light interval, metabolic flux distributions are first calculated assuming θ = 0. The ideal photon uptake rate is then multiplied by θ and set as the new upper boundary for further calculation steps.

### Nitrogen and carbon uptake kinetics

The uptake rate of nitrogen r_N_ is a function of nitrogen quota (Q_n_), growth rate μ, and extracellular nitrogen concentration (N), as proposed by Adesanya et al. [13] and shown in Eqs. (10) and (11). Other parameters include: the minimum and maximum nitrogen quotas, q_n_ and q_nm_, the maximum nitrogen uptake rate ν_nm_, and nitrogen uptake half-saturation coefficient ν_nh_.

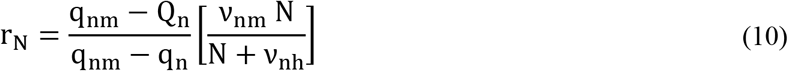

A simple mass balance on including growth-induced depletion and replenishment yields Eq. (11).

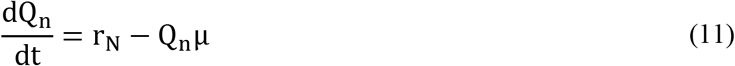

Similarly, we used the inorganic carbon uptake kinetics model proposed by Filali [11] to calculate the maximum carbon uptake rate 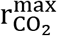 at any given moment, as shown in Eq. (12). In this model, carbon uptake is a function of the concentration of carbon dioxide C_CO_2__, the maximum carbon uptake rate from the GSM model 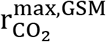, the minimum cell size Z_min_, the size increase T, biomass concentration X, and carbon uptake half-saturation coefficient K_C_.

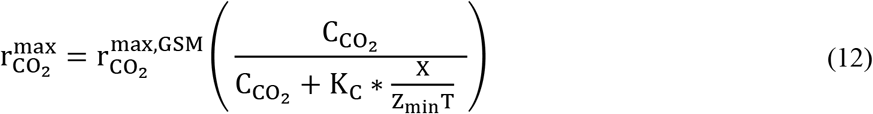

### Carbon allocation

Nutrient availability in the media directly alters the way carbon is distributed across the cell. During nutrient-sufficient conditions, microalgae tend to allocate carbon on amino acid and nucleic acid biosynthesis (herein *active biomass* or X); on the other hand, nutrient-depletion, and in general stress conditions, causes metabolism to shift carbon flow towards lipid biosynthesis. In photobioreactors, the nitrogen poses as bottleneck for overall growth, but also as trigger for lipid accumulation [6,13,17,20,51].

We proposed a simple flow distribution algorithm, with cell size (Z) and nitrogen quota (Q_n_) as coefficients for the estimation of carbon allocation. Increased nitrogen quota favored biosynthesis of active biomass and starch, whereas low nitrogen levels shifted carbon flow towards lipid production. We defined a magnitude n, which played the role of a penalty function on active biomass production and followed the Michaelis-Menten-type function shown in Eq. (13).

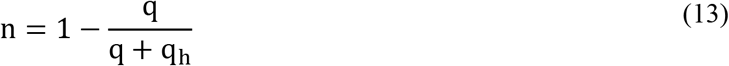

Relative intracellular nitrogen levels are represented by q = Q_n_/q_nm_, and q_h_ is the half-saturation coefficient. In a similar fashion, decreased cell sizes favored the uptake of inorganic carbon and the accumulation of storage molecules, while bigger cells were assumed to lower carbon uptake levels, as previously reported by the studies of Taguchi et al. [52] and Thompson et al. [53]. Therefore, we defined a penalty function *z* on inorganic carbon uptake, presented in Eq. (14).

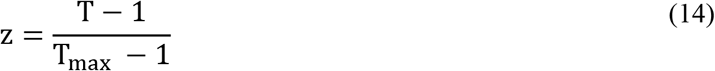

Where T is the size increase, calculated as a function of the intracellular content of starch (x_starch_) and lipids (x_lipid_), as shown in Eq. (15).

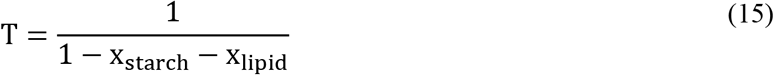

Finally, storage starch consumption is limited by a third penalty function based on the intracellular starch concentration c_starch_ and K as a half-saturation coefficient.

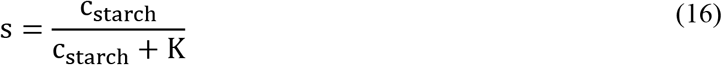

In the end, the penalty functions were used to constrain the solution space of the GSM, as upper or lower boundaries, as shown in Eq. (17) to (20). Every variable with superscript *max* is internally calculated in the algorithm as the maximum possible value at any given time point and light interval. During light and dark periods, the objective functions were, respectively, starch accumulation and biomass production, following previous reports of peak activities of starch production and consumption in light and dark periods, respectively [38–40]. An overview of the carbon allocation algorithm is illustrated in Fig S2.

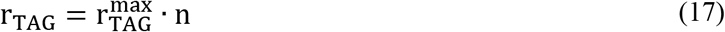

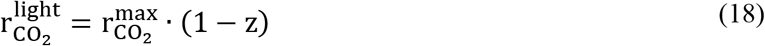

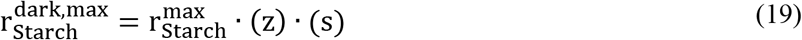

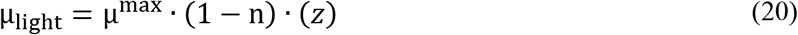

### Parameter estimation

Metabolic capabilities across species and even strains do not remain constant. This has been one of the most significant drawbacks when trying to generate a wide-spectrum biological model. However, in this work we were able to identify five strain-specific parameters which are assumed to be inherent in the microorganism: maximum size increase z_max_, maximum oxygen evolution 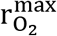 maximum carbon uptake 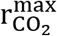, nitrogen quota half-saturation coefficient q_h_, and starch accumulation half-saturation coefficient K. Parameter estimation was done with MATLAB Optimization Toolbox, using the *active-set* algorithm. As a result, this model is capable of predicting the macroscopic outcome of a photobioreactor under different conditions for a single strain if these parameters are known. For each study we used one of the available sets of kinetic data to calculate these parameters, specifically data at an initial nitrogen concentration of 0.021 g L^−1^ for Adesanya et al. [13] and data at an irradiance of 848 μmol m^−2^ s^−1^ for Kim et al. [14]. Regression parameter values are shown in Table S1, and a summary of all other parameters is shown in Table S2.

### Maximization of lipid productivity

Five variables were manipulated to search for the optimal global lipid productivity (*R_L_* in Eq. (21)) condition: initial lamp irradiance I_0_, final lamp irradiance I_f_, photoperiod p, culture duration t_f_, and a coefficient b_I_ which represents the shape of the light profile, as shown in Eq. (22).

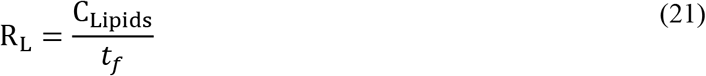

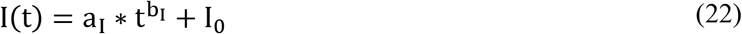

In Eq. (22), only b_I_ is used as an optimization variables, as a_I_ is dependent on the variables I_0_, I_f_ and b_I_ itself, as presented in Eq. (23).

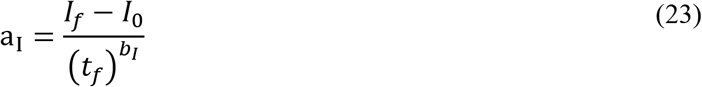

## Declarations

### Authors’ contributions

JT and RG conceived the study. JT analyzed the data; JT, CZ, JB, RG and KZ discussed the data; JT wrote the manuscript with input of all co-authors.

## Acknowledgements

JT would like to thank Ana Isabel Ramos Murillo and Johan Andrés Pasos Panqueva (UNAL-Bogota) for fruitful discussions.

## Availability of data and materials

The datasets analyzed are included in this article. The code used for simulations is fully available in the *pbr* repository at https://github.com/jdtibochab/pbr.

## Consent for publication

Not applicable.

## Ethics approval and consent to participate

Not applicable.

