## Supplementary figures and images for "A multiscale model predicts the sensitivity of *Chlorella vulgaris* to light and nitrogen levels in photobioreactors"

### Fig S1

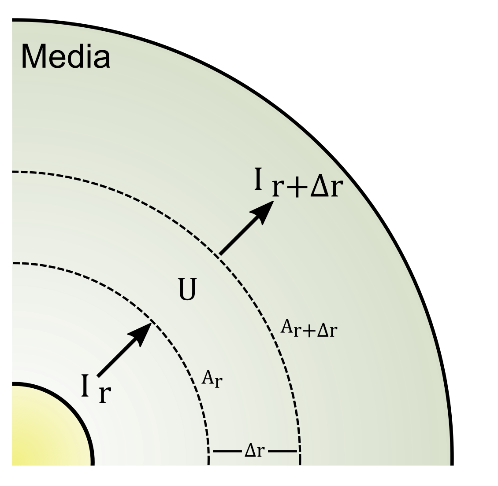

**Fig S1. Differential photon conservation balance in a photobioreactor.**

### Fig S2

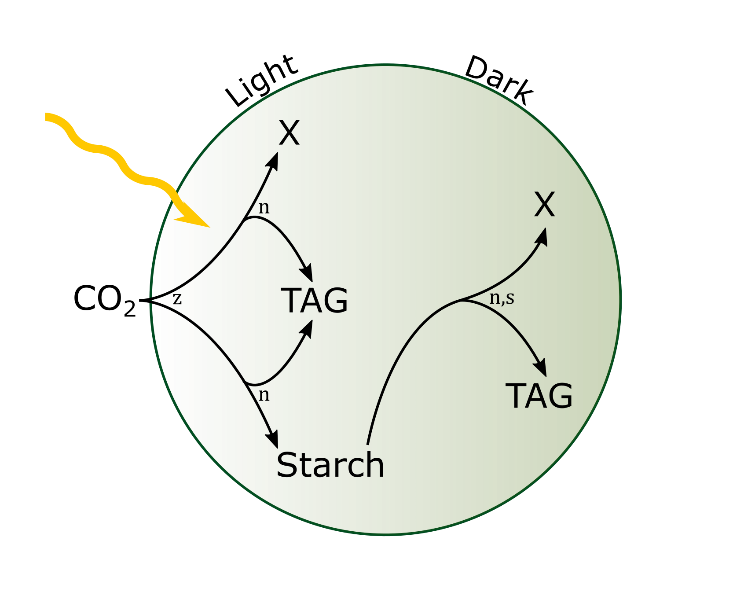

**Fig S2. Illustration of carbon allocation algorithm.**
