## Supplementary material for "A multiscale model predicts the sensitivity of *Chlorella vulgaris* to light and nitrogen levels in photobioreactors": Fig S3

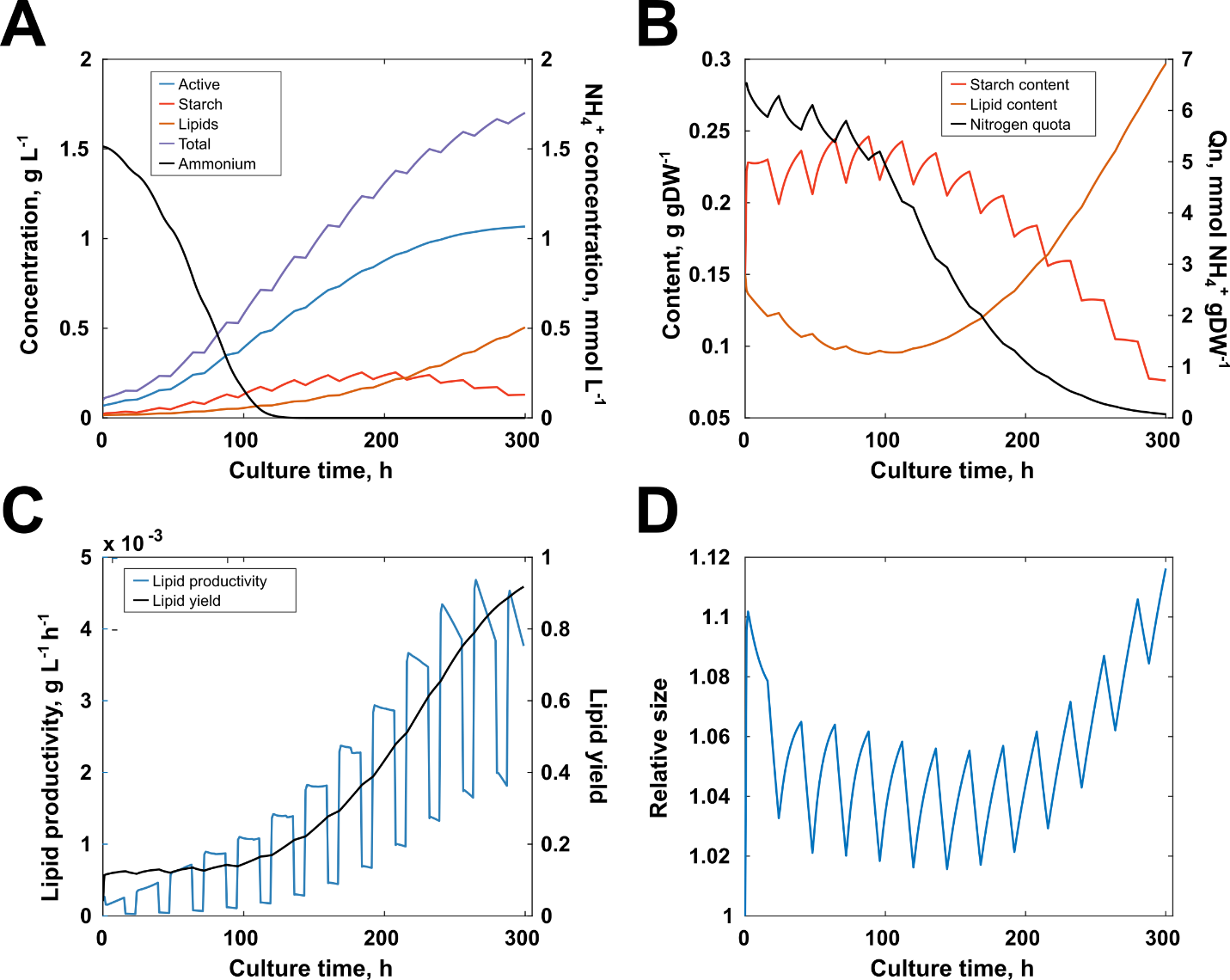

**Fig S3. Simulation results of the growth of *Chlorella vulgaris* at a hypothetical initial condition for the optimization of lipid productivity.** (A) Global reactor concentrations of active biomass, starch, lipids, total biomass and ammonium. (B) Intracellular content of starch, lipids and nitrogen. (C) Contrast of lipid productivity with lipid yield. (D) Variation of cell size.
