## Supplementary material for "A multiscale model predicts the sensitivity of *Chlorella vulgaris* to light and nitrogen levels in photobioreactors": Table S1

**Table S1. Parameter regression results for both studies included in this work.**

| **Dataset** | $\boldsymbol{z}_{\mathbf{max}}$ | $\mathbf{r}_{\mathbf{O}_{\mathbf{2}}}^{\mathbf{max}}$  $\mathbf{mmol} \mathbf{gDW}^{-\mathbf{1}}\mathbf{h}^{-\mathbf{1}}$ | $\mathbf{r}_{\mathbf{CO}_{\mathbf{2}}}^{\mathbf{max}}$  $\mathbf{mmol} \mathbf{gDW}^{-\mathbf{1}}\mathbf{h}^{-\mathbf{1}}$ | $\mathbf{q}_{\mathbf{h}}$  $\mathbf{g} \mathbf{gDW}^{-\mathbf{1}}$ | $\mathbf{K}$  $\mathbf{g} \mathbf{L}^{-\mathbf{1}}$ |
| --- | --- | --- | --- | --- | --- |
| Adesanya et al. [13] | 2.94 | 5.29 | -4.64 | 0.040 | 0.034 |
| Kim et al. [14] | 1.74 | 8.84 | -5.18 | 0.049 | 0.124 |

Five strain-specific parameters were identified to allow for the prediction of growth of a single strain under different growth conditions. These are: maximum size increase $z_{\max}$, maximum oxygen evolution $r_{O_{2}}^{\max}$, maximum CO_2_ uptake rate $r_{CO_{2}}^{\max}$, intracellular nitrogen half-saturation coefficient $q_{h}$, and starch consumption half-saturation coefficient $K$.
