## Supplementary material for "A multiscale model predicts the sensitivity of *Chlorella vulgaris* to light and nitrogen levels in photobioreactors": Table S2

**Table S2. Summary of model parameters.**

| **Parameter** | $\boldsymbol{Symbol}$ | $\boldsymbol{Definition}$ | $\boldsymbol{Units}$ | **Ref.** |
| --- | --- | --- | --- | --- |
| Light intensity profile | $I$ | $I\left( r,X \right)$ | $\mu mol m^{-2} s^{-1}$ | [22] |
| Light uptake rate | U | $U\left( r,X \right)$ | $\mathrm{mmol}\mathrm{gDW}^{-1}h^{-1}$ | $-$ |
| Fraction of active photo-system II (PSII) | $\theta$ | $\theta\left( \theta,I,t \right)$ | $-$ | [30] |
| Maximum attenuation coefficient | $K_{a,max}$ | 1041 | $m^{-1}$ | [22] |
| Light modeling parameters | $b$ | 1.03 | $\mathrm{kg}m^{-3}$ | [22] |
|  | $w$ | -0.3128 | $-$ | [22] |
|  | $p_{k}$ | 12.66 | $-$ | [22] |
| First-order PSII photoreparation coefficient | $k_{r}$ | 0.7 | $h^{-1}$ | [30] |
| First-order PSII photodamage coefficient | $k_{d}$ | $k_{d}(I)$ | $m^{2} s \mu\mathrm{mol}^{-1} h^{-1}$ | [30] |
| Photoinhibition parameters | $m_{k}$ | 0.00042 | $m^{4} s^{2} \mu\mathrm{mol}^{-2} h^{-2}$ | [30] |
|  | $b_{k}$ | 0.05 | $m^{2} s \mu\mathrm{mol}^{-1} h^{-1}$ | [30] |
| Nitrogen uptake rate | $r_{N}$ | $r_{N}(Q_{n}.N)$ | $\mathrm{mmol}\mathrm{gDW}^{-1}h^{-1}$ | [9] |
| Nitrogen quota | $Q_{n}$ | $Q_{n}\left( r_{N},Q_{n},\mu\right)$ | $\mathrm{mmol}\mathrm{gDW}^{-1}$ | [9] |
| Maximum nitrogen quota | $q_{\mathrm{nm}}$ | 6.78 | $\mathrm{mmol}\mathrm{gDW}^{-1}$ | [9] |
| Minimum nitrogen quota | $q_{n}$ | 2.29 | $\mathrm{mmol}\mathrm{gDW}^{-1}$ | [9] |
| Maximum nitrogen uptake rate | $\nu_{\mathrm{nm}}$ | 2.02 | $\mathrm{mmol}\mathrm{gDW}^{-1}h^{-1}$ | [9] |
| Nitrogen uptake half-saturation coefficient | $\nu_{\mathrm{nh}}$ | 4.29 | $\mathrm{mM}$ | [9] |
| Carbon uptake half-saturation coefficient | $K_{C}$ | 0.0128 | $\mathrm{mmolN}\mathrm{cell}^{-1}$ | [33] |
| Nitrogen-dependent penalty function | $n$ | $n\left( q \right)$ | $-$ | $-$ |
| Relative nitrogen quota | $q$ | $q\left( Q_{n} \right)$ | $-$ | $-$ |
| Size-dependent penalty function | $z$ | $z\left( Z \right)$ | $-$ | $-$ |
| Cell size | $Z$ | $Z\left( x_{\mathrm{starch}},x_{lipid} \right)$ | $pg cell^{-1}$ |  |
| Minimum cell size | $Z_{\min}$ | 75 | $pg cell^{-1}$ | [40] |
| Cell size increase | $T$ | $T(Z)$ | $-$ | $-$ |
| Intracellular starch mass fraction | $x_{\mathrm{starch}}$ | $-$ | $-$ | $-$ |
| Intracellular lipid mass fraction | $x_{\mathrm{lipid}}$ | $-$ | $-$ | $-$ |
| Starch content-dependent penalty function | $s$ | $s\left( C_{\mathrm{starch}} \right)$ | $-$ | $-$ |
| Lipid production rate | $r_{TAG}$ | $r_{TAG}(n)$ | $\mathrm{mmol}\mathrm{gDW}^{-1}h^{-1}$ | $-$ |
| Maximum CO_2_ uptake in GSM | $r_{\mathrm{CO}_{2}}^{max,GSM}$ | $-$ | $\mathrm{mmol}\mathrm{gDW}^{-1}h^{-1}$ | [3] |
| Maximum lipid production rate | $r_{\mathrm{TAG}}^{\max}$ | $-$ | $\mathrm{mmol}\mathrm{gDW}^{-1}h^{-1}$ | $-$ |
| Starch consumption rate | $r_{\mathrm{Starch}}^{dark,\max}$ | $r_{\mathrm{Starch}}^{\mathrm{dark}}(s,z)$ | $\mathrm{mmol}\mathrm{gDW}^{-1}h^{-1}$ | $-$ |
| Maximum starch production rate | $r_{\mathrm{Starch}}^{\max}$ | $-$ | $\mathrm{mmol}\mathrm{gDW}^{-1}h^{-1}$ | $-$ |
| CO_2_ consumption rate | $r_{\mathrm{CO}_{2}}^{\mathrm{light}}$ | $r_{\mathrm{CO}_{2}}^{\mathrm{light}}(s,z)$ | $\mathrm{mmol}\mathrm{gDW}^{-1}h^{-1}$ | $-$ |
| Maximum CO_2_ consumption rate | $r_{\mathrm{CO}_{2}}^{\max}$ | $-$ | $\mathrm{mmol}\mathrm{gDW}^{-1}h^{-1}$ | $-$ |
| Specific active biomass production rate | $\mu^{\mathrm{light}}$ | $\mu^{\mathrm{light}}(n,z)$ | $h^{-1}$ | $-$ |
| Global lipid productivity | $R_{L}$ | $R_{L}(C_{\mathrm{lipids}},t_{f})$ | $g L^{-1}$ | $-$ |
| Global lipid concentration | $C_{\mathrm{lipids}}$ | $-$ | $g L^{-1}$ | $-$ |
| Global CO_2_ concentration | $C_{CO_{2}}$ | $-$ | $g L^{-1}$ | $-$ |
